# A phylogenetically diverse class of “blind” type 1 opsins

**DOI:** 10.1101/033027

**Authors:** Erin A. Becker, Andrew I. Yao, Phillip M. Seitzer, Tobias Kind, Ting Wang, Rich Eigenheer, Katie S. Y. Shao, Vladimir Yarov-Yarovoy, Marc T. Facciotti

## Abstract

Opsins are photosensitive proteins catalyzing light-dependent processes across the tree of life. For both microbial (type 1) and metazoan (type 2) opsins, photosensing depends upon covalent interaction between a retinal chromophore and a conserved lysine residue. Despite recent discoveries of potential opsin homologs lacking this residue, phylogenetic dispersal and functional significance of these abnormal sequences have not yet been investigated. We report discovery of a large group of putatively non-retinal binding opsins, present in a number of fungal and microbial genomes and comprising nearly 30% of opsins in the *Halobacteriacea*, a model clade for opsin photobiology. Based on phylogenetic analyses, structural modeling, genomic context and biochemistry, we propose that these abnormal opsin homologs represent a novel family of sensory opsins which may be involved in taxis response to one or more non-light stimuli. This finding challenges current understanding of microbial opsins as a light-specific sensory family, and provides a potential analogy with the highly diverse signaling capabilities of the eukaryotic G-protein coupled receptors (GPCRs), of which metazoan type 2 opsins are a light-specific sub-clade.

## Introduction

Opsin proteins catalyze light-dependent processes in all three domains of life, including vision and circadian cycling in animals (Koyanagi & Terakita, 2013), as well as chlorophyll-independent phototrophy, osmoregulation and phototaxis in bacteria, archaea, microbial eukaryotes, and multi-cellular fungi (Inoue *et al*, 2014). The evolutionary relationship between microbial (type 1) opsins and metazoan (type 2) opsins is unresolved (Larusso *et al*, 2008; Mackin *et al*, 2014; Shen *et al*, 2013), however, in both groups, photosensing depends upon a covalent interaction between a conserved lysine residue in the seventh transmembrane helix (bovine visual rhodopsin K296 / bacteriorhodopsin K216) and a retinal chromophore (Nathans & Hogness, 1983; Mullen *et al*, 1981). Recent studies have reported discovery of genes encoding opsin homologs lacking this residue in fungal, haloarchaeal and placozoan genomes (Spudich *et al*, 2000; Siddaramappa *et al*, 2012; Feuda *et al*, 2012). However, these have been treated as isolated instances and the phylogenetic dispersal and functional significance of these abnormal sequences have not yet been investigated.

Here we report discovery of a large group of putatively non-retinal binding opsins comprising nearly 30% of opsin homologs in the archaeal family *Halobacteriacea*, a historically important model clade for study of opsin photobiology. This family of extremely halophilic archaea possesses a diverse range of opsins, which have classically been divided into four groups: the ion pumps halorhodopsin (HR) and bacteriorhodopsin (BR), which respectively regulate cytoplasmic osmolarity and create electrochemical gradients used in ATP production; and two classes of sensory rhodopsins (SR1 and SR2), which serve as histidine kinase response regulators for phototactic and photophobic behaviors (Fu *et al*, 2010). Recent studies have expanded our view of haloarchaeal opsin diversity by revealing a third sensory rhodopsin (SR3), a second group of bacteriorhodopsins (BR2), and a proposed intermediate between bacteriorhodopsin and the sensory rhodopsins (MR) (Baliga *et al*, 2004; Fu *et al*, 2010; Bolhuis *et al*, 2006; Sudo *et al*, 2011). Studies of these diverse haloarchaeal opsins have lead to major advances in our understanding of the kinetics and structural intermediates of opsin photocycles (Landau *et al*, 2003), spectral tuning (Wang *et al*, 2014), and signal transduction pathways (Grote *et al*, 2014). Haloarchaeal opsins have also served as models for protein crystallization (Grote *et al*, 2014), membrane protein folding (Tastan *et al*, 2014) and development of optogenetic toolkits (Zhang *et al*, 2011).

Based on genomic context, phylogenetic analyses, structural modeling and biochemistry, we propose that these abnormal opsin homologs, which are also present in some fungal, cyanobacterial, and chlorophytal genomes, represent a novel family of sensory opsins potentially involved in taxis response to one or more non-light stimuli. This finding challenges current understanding of microbial type 1 opsins as a light-specific sensory family, and provides a potential analogy with the highly diverse signaling capabilities of the eukaryotic G-protein coupled receptors (GPCRs), of which type 2 opsins are a light-specific sub-clade. These results call for a renewed perspective on the roles of type 1 opsins in microbial physiological responses to diverse environmental inputs.

## Results and Discussion

A large-scale survey of 80 complete and high-quality draft haloarchaeal genomes (Becker *et al*, 2014) revealed a novel, large opsin class, consisting of 48 homologs lacking the normally conserved lysine residue (K216) required for binding retinal out of 170 total haloarchaeal opsins (Fig 1, [Data Dryad link]). This group of sequences, hereafter termed the retinal-free opsins (RFOs) thus comprises nearly 30% of all known haloarchaeal opsins. We deliberately use the term opsin to describe the RFOs, even though the term is traditionally used to describe the apoprotein of the retinal-bound rhodopsin, due to their proposed constitutively retinal-free nature and phylogenetically close relationship to sensory opsins. The RFOs are broadly distributed across 11 genera in all three major haloarchaeal clades (Becker *et al*, 2014), in species with and without canonical opsin homologs (Table S1). All eight species whose genomes encode RFOs and lack canonical opsin homologs were also found to lack *crtY* and *brp*, genes encoding enzymes which catalyze the terminal steps in retinal biosynthesis (McCarren & DeLong, 2007). Together with additional biochemical and structural modeling, these data suggest the hypothesis that RFO genes encode type 1 opsins that have a non-retinal dependent function, providing a potential functional analogy with eukaryotic GPCRs.

**Figure 1:**
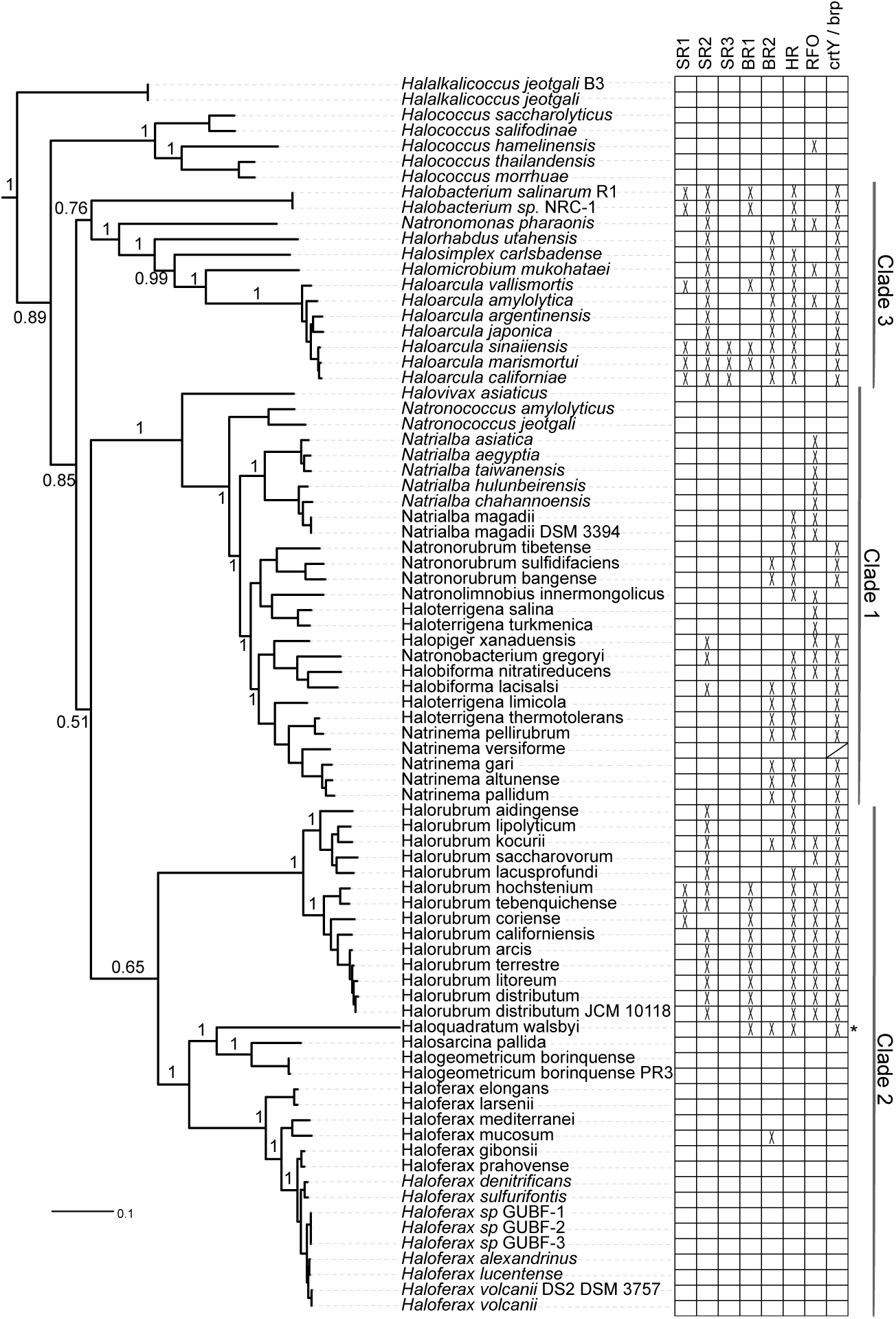
Phylogenetic distribution of opsin classes across the Haloarchaea. Distribution of six previously characterized haloarchaeal opsin families, the putative retinal-free opsins, and the retinal biosynthesis genes *crtY* and *brp* are shown superimposed on a multi-marker phylogenetic tree of 80 sequenced haloarchaea (tree published in Becker *et al*, 2014). SR = sensory rhodopsin, BR = bacteriorhodopsin, HR = halorhodopsin, RFO = retinal-free opsin. Asterisk indicates the presence of middle rhodopsin (MR) (Sudo *et al*, 2011). Phylogenetic distribution of *crtY* and *brp* are identical except for one species (*Natrinema versiforme*) which has *brp* but no detected *crtY* homolog (marked with /). Haloarchaeal clade designations as in Becker *et al*, 2014. Bootstrap support values for lower-level clades removed for clarity. Tree file can be accessed at [Data Dryad link].

Phylogenetic analysis revealed that the RFOs form a monophyletic clade most closely related to sensory opsins (Fig 2A) and themselves sub-divide into two distinct groups, one consisting of sequences primarily from *Halorubrum* species (group A) and the other of sequences from *Natrialba* species and other Clade 1 haloarchaea (group B) (Fig 2B). In group A RFOs, the Schiff base lysine was replaced with arginine, while in group B RFOs this position contained a leucine or other hydrophobic residue. All but one of the 16 group A RFOs were located adjacent to a predicted methyl-accepting chemotaxis (HAMP/MCP) signal transducer, providing evidence for the hypothesis that the RFOs represent a novel form of sensory opsins, which are often co-operonic with their cognate signal transducers (Spudich, 2006) (Fig. S1A). Many (17/32) group B RFOs were similarly linked to HAMP/MCP family signal transducers, with nine also located in the proximity of chemotaxis and flagellar biosynthesis operons (Fig. S1B). This close genomic association suggests a functional role for at least a sub-set of RFOs in modulating the flagellar apparatus in response to an as yet un-identified signal(s). The remaining 15 group B RFOs, not located adjacent to signal transducer genes, belonged to 10 species, three of which have been described as non-motile (Xu *et al*, 1999, 2001). We therefore propose that a number of group B RFOs may have signal-response functions unrelated to motility, as is the case for a large number of GPCR homologs (Schiöth & Fredriksson, 2005) and has been suggested for *Anabaena* sensory rhodopsins (ASRs) (Brown, 2014). Several of the RFOs not linked to signal transducers were located near genes implicated in various stress responses, including heat shock proteins, metal chaperones, and carbon starvation proteins – proposing future lines of research for deciphering the functions of these group B RFOs.

**Figure 2:**
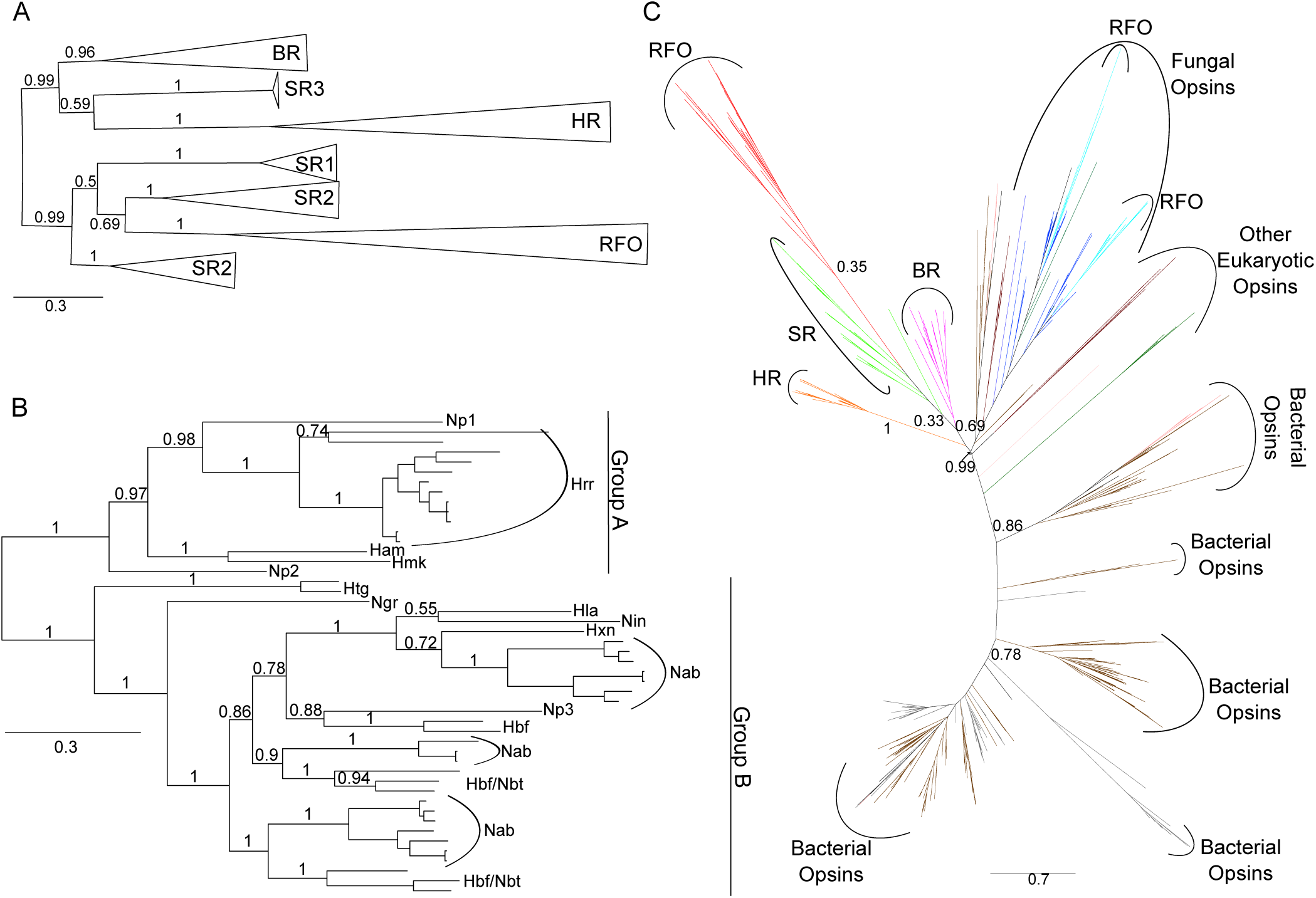
Phylogenies of haloarchaeal and microbial (type 1) opsins. **A** Phylogenetic tree of 170 haloarchaeal opsin proteins constructed using Bayesian inference with MrBayes. Individual members of each opsin family were collapsed to indicate relationship among classes. Triangle length is proportional to sequence diversity within clade, triangle width is not significant. Abbreviations as in Fig. 1. For fully expanded tree see Fig. S5. Tree file can be accessed at [Data Dryad link]. **B** Bayesian inference phylogenetic tree of novel class of putatively retinal-free sensory opsins extracted from tree in Fig. 2A. Np = *Nmn. pharaonis*, Hrr = *Halorubrum*, Ham = *Har. amylolytica*, Hmk = *Hmc. mukohataei*, Htg = *Haloterrigena*, Ngr = *Nbt. gregoryi*, Hla = *Hbf. lacisalsi*, Nin = *Nln. innermongolicus*, Hxn = *Hpg. xanaduensis*, Nab = *Natrialba*, Hbf = *Halobiforma*, Nbt = *Natronobacterium*. **C** Maximum-likelihood phylogeny of all microbial (type 1) opsins obtained by BLASTp search of NCBI’s nr and env_nr databases. Tree inferred using FastTree (Price *et al*, 2010) and ComparetoBootstrap.pl (Price) using 500 bootstrap replicates generated with SeqBoot (Felsenstein, 2005). Branches are colored by phylogenetic affiliation and bootstrap support values above 0.30 are shown for major clades. Colors: grey = unannotated/unassigned, brown = bacterial, salmon = dinoflagellates, dark green = viridiplantae, purple = haloarchaeal BR, light green = haloarchaeal SR, orange = haloarchaeal HR, red = haloarchaeal RFO, light blue = fungal RFO, dark blue = other fungal opsin. Abbreviations as in Fig. 1. For fully expanded tree see Fig. S3. Tree file can be accessed at [Data Dryad link].

Both haloarchaeal bacteriorhodopsin and bovine visual rhodopsin have been shown to function at reduced efficiencies in the absence of K216/K296, when aminated retinylidene compounds are provided in lieu of retinal (Zhukovsky *et al*, 1991; Schweiger *et al*, 1994; Friedman *et al*, 1994). To investigate the possibility that the RFOs may function in a canonical light-sensitive manner by retaining interaction with retinal, or a retinal-like chromophore, but not Schiff base formation at K216, we conducted residue conservation analysis and structural modeling. Of three residues experimentally characterized as providing a hydrophobic cavity for the retinal ring in *Natronomonas pharaonis* SR2 (V108, F127, W178) (Pebay-Peyroula *et al*, 2002), only one is conserved as a hydrophobic residue of similar size in group A RFOs (M108) and none in group B RFOs (Fig 3A). W178, which is universally conserved as an aromatic residue in canonical haloarchaeal opsins, has been converted to the much smaller alanine and glycine residues in group A RFOs and group B RFOs, respectively. Similarly, the highly conserved aromatic residue at position 127 has been converted to a polar amino acid (T/S) in most RFOs. In addition, group A RFOs are missing two (W76 ➔ D, Y174 ➔ L) and group B RFOs one (Y174 ➔ L) of the aromatic residues involved in steric constraint of the retinal polyene chain (Pebay-Peyroula *et al*, 2002) (Fig 3A). These results strongly suggest that, in addition to having lost the Schiff base lysine for covalent binding of retinal, RFOs lack the canonical binding pocket to accommodate retinal-like chromophores. Conversely, residues involved in sensory signaling are conserved in one or both RFO clades. Y51 and R72, which together form a water-mediated hydrogen-bond complex important in propagation of signal to the linked signal transducer (Ishchenko *et al*, 2013), are highly conserved in group B, but not group A RFOs. Y199, which forms a hydrogen bond with the signal transducer in *Natronomonas pharaonis* SR2 (Gordeliy *et al*, 2002), is universally conserved across the RFO clade. Similarly, D189, which in *Nmn. pharaonis* SR2 also hydrogen bonds with the cognate signal transducer (Gordeliy *et al*, 2002), is highly conserved in both group A and group B RFOs, however, this residue is poorly conserved in other canonical SR2 homologs. The high level of conservation of residues involved in signal transduction, combined with lack of conservation of the retinal binding pocket, strongly suggest that RFOs are a non-retinal utilizing family of sensory opsins.

**Figure 3:**
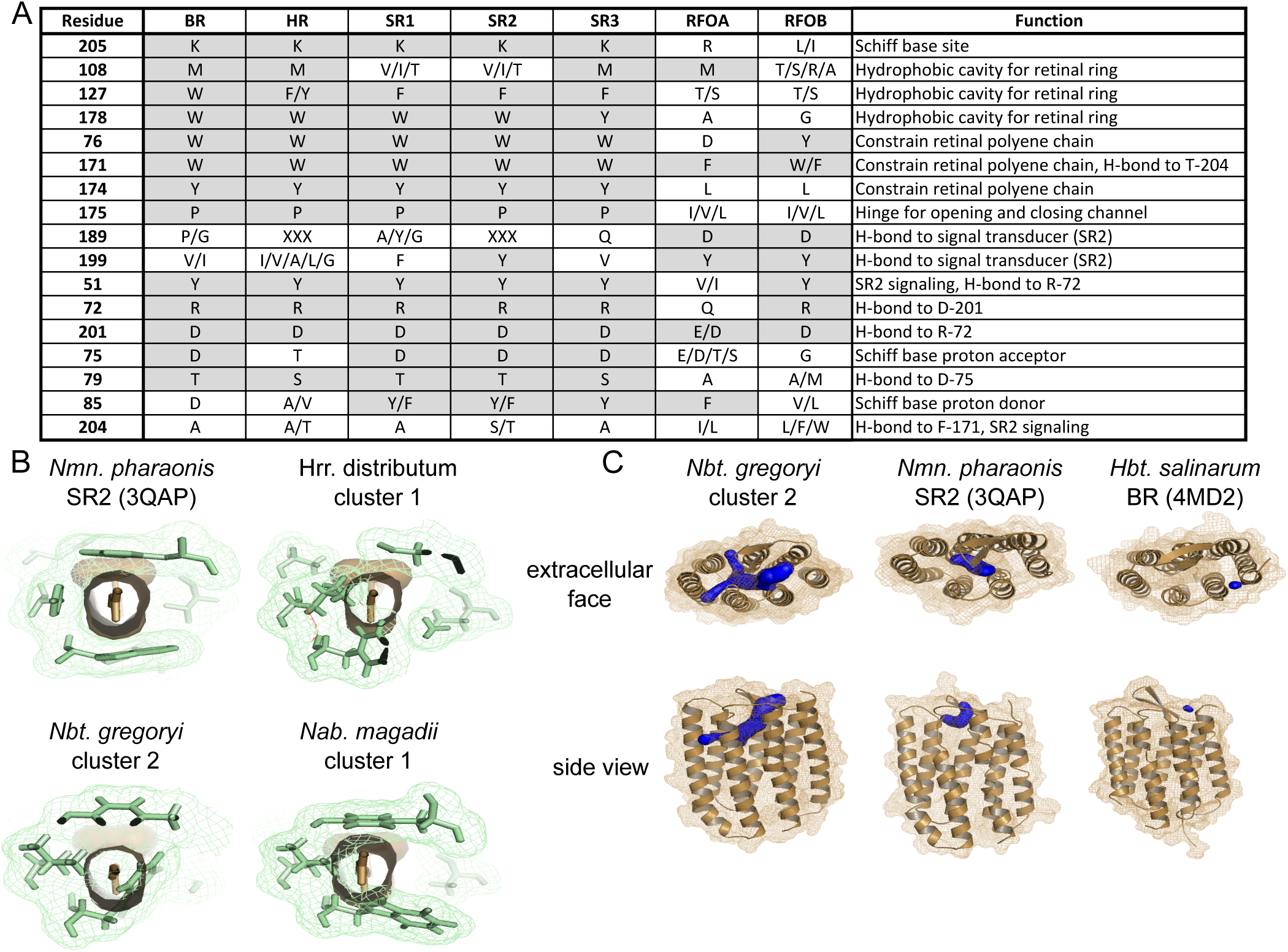
Residues important for signal propagation, but not retinal binding, are conserved across haloarchaeal RFO homologs. **A** Conservation patterns in positions of functional importance across five canonical opsin and two RFO subclasses show that residues involved in signaling, but not those involved in retinal binding, are conserved in RFOs. XXX = highly variable. Positions with conserved or biochemically similar amino acid residue across multiple opsin classes are shaded. Residue numbering is according to *Nmn. pharaonis* SR2 (YP_331142). **B** Residues forming the retinal binding cavity in canonical opsins show steric clash with canonical retinal binding pocket in predicted RFO structural models. Residues corresponding to *Nmn. pharaonis* SR2 V108, F127, W178, W76, W171, and Y174 are shown in green with lattice representing atomic surface. Retinal is shown in tan, looking down polyene chain, with solid tan representing atomic surface. **C** Representative output visualizations from CAVER (Chovancova *et al*, 2012) showing predicted novel extracellularly accessible binding pocket and internal tunnel network for *Nbt. gregoryi* WP_005575895. Similar search parameters revealed much smaller (SR2, BR) or no (HR) cavities for canonical opsins. For all CAVER predictions, see Fig. S2.

Structural models of three RFO homologs, using *Natronomonas pharaonis* SR2 (PDB ID: 3QAP) as a template, were consistent with residue analysis in showing lack of a canonical retinal binding pocket. Residues homologous to those lining the canonical binding pocket were shown to overlap the space normally occupied by retinal (Fig 3B). Additionally, a novel extracellularly accessible binding pocket was identified which was missing or highly diminished in canonical haloarchaeal opsins (Fig 3C, Fig. S2). To identify candidate ligands for these novel binding pockets, we used the program idock (Li *et al*, 2012) to screen 229,358 natural product ligands against the three structurally modeled RFO homologs. An overlap analysis of the results showed that, despite their structural similarity, the three opsins likely have affinities for very different compounds. Nevertheless, most were sourced from the same ligand classes with similar scaffolds including oxygen and nitrogen heterocycles and sesquiterpene (See SI Discussion). Many examples of extracellularly accessible ligand binding pockets exist for Class A (rhodopsin-like) GPCRs (Tan *et al*, 2013; Haga *et al*, 2012; Rasmussen *et al*, 2011; Lebon *et al*, 2011), further suggesting a non-light related signal response function for the RFOs. Although both structural modeling and residue analysis suggest that the RFOs lack retinal binding capabilities, the possibility that they may utilize an alternate chromophore remains to be tested.

As type 1 opsin homologs lacking the Schiff base lysine have previously been reported for fungi (Spudich *et al*, 2000), we were interested in determining whether haloarchaeal and fungal RFOs are monophyletic or represent independent losses of retinal-binding ability. We collected type 1 opsin homologs from the NBCI’s nr and env_nr databases and performed maximum-likelihood phylogenetic analysis. A total of 1,077 opsin sequences were included. Interrogation of alignments for sequences lacking the Schiff base lysine revealed 45 non-haloarchaeal RFOs, none of which branched within the haloarchaeal RFO clade (Fig 2C). Thus, the haloarchaeal RFOs are evolutionarily distinct from other potentially non-light sensitive microbial opsins. The 45 non-haloarchaeal RFOs included two groups of fungal homologs, one containing 31 sequences from 10 Ascomycota genera and the other eight sequences from seven genera spanning the Ascomycota and Basidomycota. The remaining six sequences were singletons scattered across the phylogeny (Fig. S3). Thus, although loss of the Schiff base lysine, and therefore probable loss of retinal-binding ability, has occurred at least nine times in the evolutionary history of type 1 microbial opsins, the expansion and diversification of both haloarchaeal and fungal RFOs make these exciting targets for learning novel signal transduction strategies used by type 1 opsins.

We also performed a targeted screen for haloarchaeal-type RFOs in the NCBI nr and env_nr databases, as well as the CAMERA metagenomic databases. We detected only one new non-haloarchaeal RFO belonging to the basidiomycete *Trametes versicolor* (EIW60452), which also possesses a fungal-type RFO (EIW51701). The haloarchaeal-type *Trametes versicolor* RFO branched with haloarchaeal ion pumps (BR/HR) rather than the haloarchaeal RFOs and SRs (Fig. S4). We therefore propose that this sequence represents horizontal gene transfer of either BR or HR from the Haloarchaea, followed by loss-of-function, rather than fungal acquisition of a haloarchaeal RFO homolog.

As all previous mentions of K216/K296-lacking opsins in the literature have been based on genomic data (Spudich *et al*, 2000; Siddaramappa *et al*, 2012; Feuda *et al*, 2012), and these sequences have not yet been shown to be expressed, here we verified transcription of several RFO homologs in four haloarchaeal species *(Halorubrum litoreum* JCM 13561, *Halorubrum distributum* JCM 9910, *Natrialba magadii* DSM 3394 and *Natronobacterium gregoryi* SP2). Eight of the nine investigated RFO homologs were shown to be transcribed under standard laboratory conditions (Fig 4A). For one species *(Hrr. distributum)*, transcription was also interrogated under conditions of maximal salt tolerance (5.0 M NaCl), reduced dissolved oxygen (shaking at 50 rpm vs 350 rpm), and high cell density (stationary phase). RFO transcript was detected under all conditions. Despite robust transcriptional response, we were unable to detect RFO protein in native hosts by LC-MS/MS, suggesting RFO expression may be very low or under post-transcriptional control (see SI Methods). Additional work will be required to determine biologically relevant expression conditions for RFO proteins. To experimentally verify the proposed inability of RFO proteins to bind retinal, we heterologously expressed His-tagged RFOs from two species *(Hrr. distributum* and *Nab. magadii)* (Fig 4B), and incubated purified RFOs with free all-trans retinal. Neither RFO showed absorption in the 480–580 nm range characteristic of canonical opsins (Fu *et al*, 2010). This evidence further suggests that RFO proteins do not bind the retinal chromophore (Fig 4C).

**Figure 4:**
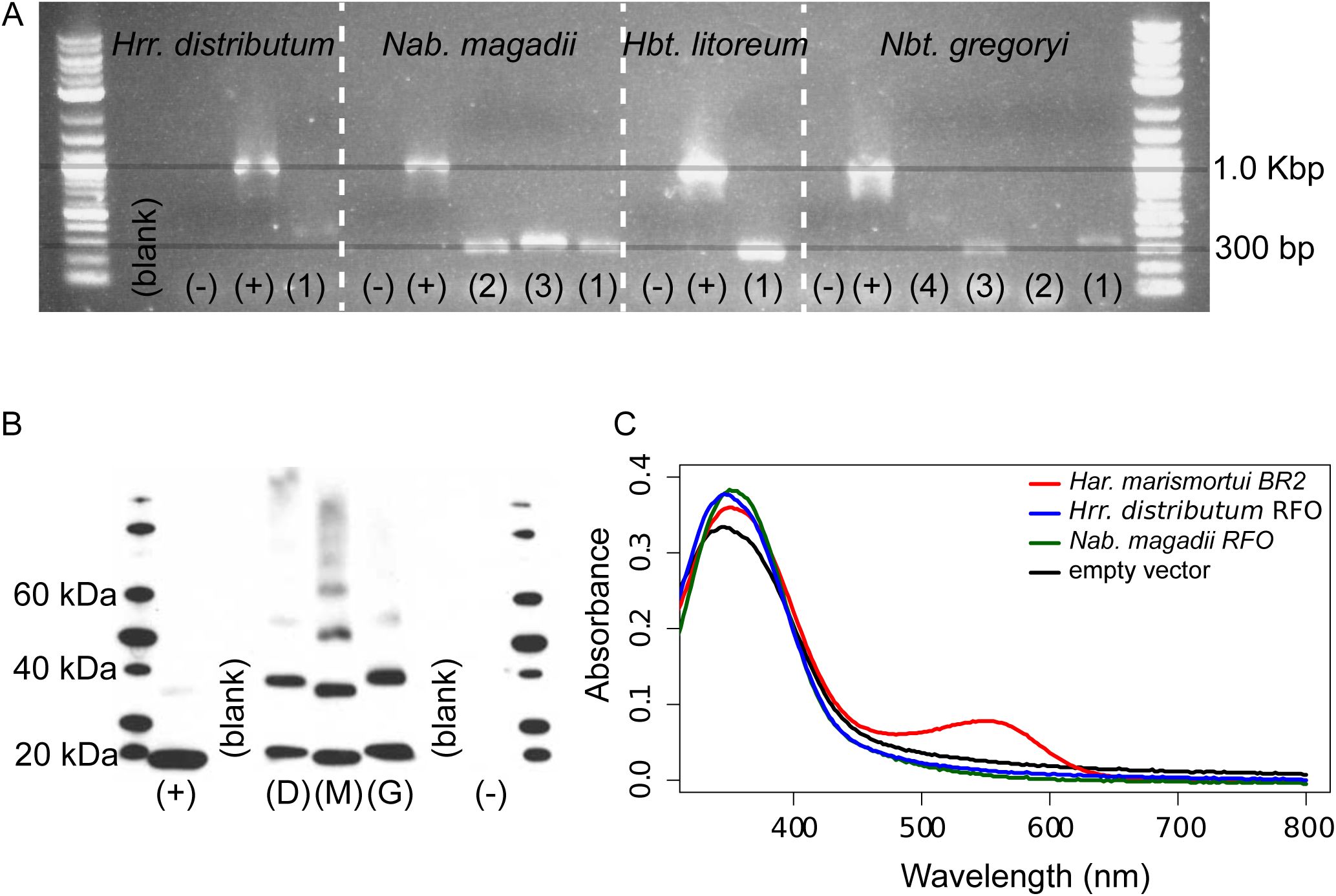
Heterologously expressed RFO homologs do not display retinal-dependent light absorption. **A** Transcription of RFOs under standard laboratory conditions was confirmed in four native hosts from both RFO clades. Eight of nine tested RFOs were transcribed, many at low levels. Positive controls are 16S rRNA gene products. *Hrr. distributum* JCM 9100 (1) = ELZ45759. *Nab. magadii* DSM 3394 (1) = WP_004215682, (2) = WP_004267173, (3) = WP_004267171. *Hbt. litoreum* (1) = WP_008366300. *Nbt. gregoryi* (1) = WP_005581268, (2) = WP_005575895, (3) = WP_015233632, (4) = WP_005579638. **B** Heterologous expression of RFO proteins was confirmed by Western blot. D = *Hrr. distributum* JCM 9100 ELZ45759, M = *Nab. magadii* DSM 3394 WP_004267173, G = *Nbt. gregoryi* DSM 3393 WP_005575895. Positive control = *Har. marismortui* BR2 YP_137573. **C** Spectra showing lack of absorption in canonical range for haloarchaeal opsins (480–580 nm) by heterologously expressed RFO homologs incubated with free-retinal. *Har. marismortui* BR2 = YP_137573, *Hrr. distributum* RFO = ELZ45759. *Nab. magadii* RFO = WP_004267173.

## Conclusions

In summary, evidence from genomic context, phylogenetic analysis, structural modeling, and biochemistry provide strong support for the existence of a large family of non-retinal binding opsins derived from a haloarchaeal-specific duplication and divergence of canonical sensory opsins, many of which are possibly involved in chemotaxis and/or transcriptional response to environmental stresses. This new family comprises nearly 30% of all known haloarchaeal opsins, providing a rich set of models through which to explore evolutionary diversification of signaling protein inputs as well as an enriched understanding of the roles played by microbial opsins in integrating diverse environmental inputs into a coordinated physiological response.

## Methods and Materials

### Haloarchaeal opsin sequence acquisition, alignment, and phylogenetic inference

During automated annotation of 80 haloarchaeal genomes with the Rapid Annotation Using Subsystem Technology (RAST) server (Aziz *et al*, 2008), five sequences were annotated as opsin homologs which did not show significant sequence similarity to canonical haloarchaeal opsins by BLASTp search (e-value cutoff of 10^−5^) and which were missing the Schiff base lysine required for binding of the retinal chromophore in all experimentally characterized opsins (K216 in the haloarchaeal model opsin *Halobacterium sp*. NRC-1 bacteriorhodopsin). These sequences were also annotated as opsins in the NCBI genomic database. A BLASTp search (e-value cutoff of 10^−5^) against the 80 haloarchaeal genomes using these five opsin sequences as queries, recovered 43 additional K216-lacking, putative opsin homologs for a total of 48 homologs in 28 genomes. These were combined with 122 canonical haloarchaeal opsins recovered by BLASTp search (e-value cutoff of 10^−5^) of the 80 haloarchaeal genomes using all *Haloarcula marismortui* ATCC 43049 opsins as query sequences. This set of queries was chosen because *Har. marismortui* possesses homologs for each of the six previously categorized classes of haloarchaeal opsins (Baliga *et al*, 2004; Fu *et al*, 2010). A total of 170 haloarchaeal opsin homologs were included in subsequent analyses ([Data Dryad link]).

Multiple sequence alignments were created separately for canonical and non-canonical haloarchaeal opsins using MUSCLE 3.8 (Edgar, 2004b, 2004a), and checked for accuracy against a previously published haloarchaeal opsin alignment (Pebay-Peyroula *et al*, 2002). Miscalled start sites for 56 sequences were manually corrected based upon alignment with sequences of experimentally characterized opsins. Individual alignments were combined using the profile-profile alignment option in MUSCLE 3.8 and manually trimmed in the alignment editor Jalview (Clamp *et al*, 2004). A total of 204 positions were used to infer phylogeny using the Bayesian tree-building software MrBayes (Huelsenbeck & Ronquist, 2001; Ronquist & Huelsenbeck, 2003) with 1.2 million rounds of Markov chain Monte Carlo iteration. The final potential scale reduction factor was 1.000 and final standard deviation of split frequencies was 0.010270. The tree was visualized using FigTree (Rambaut, 2012). The tree file is available at [Data Dryad link]. For a fully expanded tree, see Fig. S5. The alignment used for tree inference is available at [Data Dryad link].

### Distribution of opsin classes and retinal biosynthesis genes in the Haloarchaea

The presence/absence pattern of haloarchaeal opsin subclasses and the retinal biosynthesis genes *crtY, crtE, crtB, crtI*, and *brp* were superimposed on a previously published multi-marker phylogeny of the haloarchaea (Becker *et al*, 2014) using iTOL (Letunic & Bork, 2011, 2007). Subclass membership for each opsin homolog was determined based on clade affiliation in haloarchaeal opsins tree. Sequences for *crtY, crtE, crtB, crtI*, and *brp* homologs were retrieved via BLASTp searches against the local haloarchaeal database, using query sequences from six haloarchaeal species ([Data Dryad link]) and an e-value cutoff of 10^−20^. As all species with *crtY* and *brp* also had a full complement of *crtE, crtB*, and *crtI*, presence or absence of *crtY* and *brp* was used to represent retinal biosynthesis ability.

### Genome context of RFOs

Genomic context of haloarchaeal RFOs was investigated using JContextExplorer (Seitzer *et al*, 2013). Haloarchaeal genomes can be loaded using the “Retrieve Popular Genome Set” function. Annotations associated with this genome set were done using the RAST annotation service (Aziz *et al*, 2008) (see Becker *et al*, 2014). To view RFO contexts, search by Cluster Number for “4077;15722” (group A) or “2453;13537” (group B). Cluster numbers represent homology families as defined in Becker *et al*, 2014).

### Structural modeling of RFO proteins

We used x-ray structure of *Natronomonas pharaonis* SR2 (PDB ID: 3QAP) (Gushchin *et al*, 2011) as a template for generating structural models for three RFO proteins *(Nab. magadii* WP_004267173, *Hrr. distributum* ELZ45759, and *Nbt. gregoryi* WP_005575895) using the Rosetta-Membrane method (Yarov-Yarovoy *et al*, 2012, 2006; Barth *et al*, 2007). We used *Nmn. pharaonis* SR2 because it was identified by the HHpred server (Hildebrand *et al*, 2009; Söding *et al*, 2005) as the closest structural homolog to the RFOs (21–25% sequence identity). Due to a two residue deletion in the loop between TM6 and TM7 in the *Hrr. distributum* RFO compared with *Nmn. pharaonis* SR2, we predicted the structure of this region *de novo* using Rosetta cyclic coordinate descent (CCD) and kinematic (KIC) closure loop modeling, developed to model loop structures with sub-angstrom accuracy (Wang *et al*, 2007; Mandell *et al*, 2009). Several rounds of CCD and KIC loop modeling were performed with at least 10,000 models generated during each round. Models were ranked based on total Rosetta energy after each loop modeling round (Yarov-Yarovoy *et al*, 2012; Barth *et al*, 2007; Rohl *et al*, 2004). Ten percent of the lowest energy models were clustered (Bonneau *et al*, 2002) using a root mean square deviation (RMSD) threshold that placed 1–2% of all models in at least one of the largest clusters. Models representing centers of the top 20 clusters (early rounds) and/or the best 10 models by total energy (later rounds) were used as input for the next round of loop modeling. Rosetta’s full atom relaxation protocol (Barth *et al*, 2007; Rohl *et al*, 2004) was used to explore potential differences in backbone and side chain conformations of RFOs compared with the SR2 template. Selection of the best RFO models was guided by clustering of the lowest energy models to generate the most frequently sampled conformations.

### Intramolecular pathway and ligand-binding pocket predictions

Comparative structural models described above were used for prediction of ligand binding sites, internal cavities and tunnels using CAVER 3.0 (Chovancova *et al*, 2012). Analysis was performed individually for each of the top five models generated by Rosetta for each RFO, which had a mean pair-wise RMSD across all models ranging from 0.610 to 0.694 Å. CAVER settings used were minimum probe radius of 0.9 Å, shell depth of 4 Å, shell radius of 3 Å, clustering threshold of 3.5 Å, 12 approximating balls, and maximum distance and desired radius for starting point optimization of 3 Å and 5 Å, respectively. Starting point coordinates were optimized to enable use of homologous starting positions for the majority of models. The starting point for all but two of the 15 RFO models was G193 *(Nab. magadii)* / A185 *(Hrr. distributum)* / G196 *(Nbt. gregoryi)*. For two *Hrr. distributum* models, this starting position resulted in no predicted cavities, however, cavities similar to those predicted in other models were discovered with a starting position of Q175-I176. This starting position also resulted in no predicted cavities for SR2, as expected. For comparative visualization purposes, the starting position of P183 was used for SR2, resulting in prediction of a small, likely artifactual ligand binding pocket. For comparison, cavities were also predicted for bacteriorhodopsin (PMID: 4MD2) (Borshchevskiy *et al*, 2014) and halorhodopsin (PMID: 1E12) (Kolbe, 2000) homologs. Starting points for these predictions were P4 and N3, respectively. Binding pockets and tunnels were visualized with MacPyMOL (MacPyMOL). For comparison of cavity predictions between RFOs and canonical opsins, see Fig 3C. For comparison of all RFO models, see Fig. S2.

### Virtual ligand screening methods

Autodock VINA (Trott & Olson, 2010) (1.1.2 May 11, 2011), downloaded from http://vina.scripps.edu/, was used for fast protein-ligand screening. For fast multithreaded ligand screening the program idock (Li *et al*, 2012) (v2.1.3) was downloaded from https://github.com/HongjianLi/idock. For additional docking experiments we used Autodock (v4.2.6) and Autodock tools (Morris *et al*, 2009), the PyRx (Dallakyan & Olson, 2015) virtual screening software (v0.8) from http://pyrx.sourceforge.net/ as well as UCSF Chimera (Pettersen *et al*, 2004) for ligand visualization and protein modifications. For SDF file and metadata handling we used the freely available OSIRIS DataWarrior (Sander *et al*, 2015) www.openmolecules.org/datawarrior/. The statistical analysis of docking results was performed with StatSoft Statistica (v12) and computational compound clustering of results including 3D mapping was performed with CheS-Mapper (Gütlein *et al*, 2014). Venn diagrams for compound overlap based on compound ID were calculated on the web page http://www.bioinformatics.lu/venn.php.

Protein data included three canonical opsins (PDB ID: 1E12, 3QAP, and 4MD2) and three RFOs *(Nab. magadii* WP_004267173, *Hrr. distributum* ELZ45759, and *Nbt. gregoryi* WP_005575895). Receptor preparation included removal of water and unwanted ligands as well as pdbqt conversion with AutoDock tools and PyRx software.

A total of 229,358 ligands were sourced from the Universal Natural Products Database (UNPD, pkuxxj.pku.edu.cn/UNPD/) (Gu *et al*, 2013). The 3D-conformers were created using the ChemAxon molconvert software (ChemAxon molconvert; For 3D conformer generation; Molecule File Converter, version 6.0.5, 2013 ChemAxon Ltd. ChemAxon), utilizing the MMFF94 force field optimization with the lowest energy conformer option and explicit hydrogens. The SDF file was then transformed with OpenBabel (O’Boyle *et al*, 2011) (v2.3.2) into a ligand pdbqt file and subsequently split into multiple single files with VINA_split.exe. Idock configurations files were created for each receptor with a box size of 15 Å on each axis, including the following docking parameters, (x:16.560 y:40.852 z:-4.072) for *Nab. magadii* WP_004267173, (x:14.244 y: 40.502 z:-1.779) for *Hrr. distributum* ELZ45759 and (x:14.527 y:41.251 z:-2.290) for *Nbt. gregoryi* WP_005575895. All 229,358 ligands were screened for each protein. Because the scoring functions of VINA and idock are non-deterministic we repeated the scoring for the best candidates in order to screen for outliers. For the creation of a small decoy library (Mysinger *et al*, 2012), known ligands from the canonical opsins were submitted to http://dude.docking.org/generate. The decoy library with 750 compounds was then used to generate energy cutoff values for the idock-based screening.

The hardware for virtual screening included a Dual Xeon High Performance Workstation with two Intel E5–2687W processors (16 cores, 32 threads), 8 Tb RAID10 disks, 2 Tb SSDs and 196 Gb RAM under Windows 7 64-bit. All modeling was performed on an 80 Gb Soft Perfect RamDisk allowing sequential read/write speeds of up to 4 Gb/second. For additional discussion of ligand screening, see SI Discussion.

### Type 1 opsin sequence acquisition, alignment, and phylogenetic inference

Sequences for bacterial, archaeal, and microbial eukaryotic type 1 opsins were obtained by BLASTp search of the NCBI nr and env_nr databases (as of October 30^th^, 2012) with an e-value cutoff of 10^−5^ and maximum target sequences set to 100,000 using all *Har. marismortui* opsins as query sequences as described above. A total of 1,430 sequences were recovered. After removing mutants, synthetic constructs, highly fragmentary sequences and sequences lacking a predicted Bac_rhodopsin domain (Pfam clan CL0192), 907 opsin homologs remained. Sequences acquired from each database were independently aligned using MUSCLE 3.8.31 (Edgar, 2004a, 2004b), manually trimmed to remove poorly aligned regions and to account for the fragmentary nature of metagenomic sequencing, and alignments combined using the profile-profile alignment option. After adding the 170 haloarchaeal sequences from our dataset, a total of 147 positions for 1,077 opsin sequences were used in tree inference. For these sequences, an initial guide tree was constructed using FastTree (Price *et al*, 2010, 2009), then 1000 re-sampled alignments were generated using the Phylip package SEQBOOT (Felsenstein, 2005) and used to determine bootstrap support values for clades in the initial tree with CompareToBootstrap.pl (Price). The resulting tree was visualized with FigTree (Rambaut, 2012). The tree file is available at [Data Dryad link]. For fully expanded tree, see Fig. S3 (red leaf labels represent sequences lacking the Schiff base lysine).

### Phylogenetic distribution of haloarchaeal-type RFOs

All protein and peptide databases in CAMERA and the NCBI nr and env_nr databases (as of December 18^th^, 2012) were interrogated for non-canonical opsin homologs using BLASTp searches with all 48 haloarchaeal RFOs as queries. BLASTp parameters and database details are listed in Table S2. Redundant sequences were removed using Jalview (Clamp *et al*, 2004) and unique sequences searched against the Pfam database (Finn *et al*, 2010) for presence of a Bac_rhodopsin domain (CL0192). Sequences with a Bac_rhodopsin domain were aligned using MUSCLE 3.8 (Edgar, 2004a, 2004b) and presence or absence of K216 was recorded. To assess the evolutionary origin of the only non-haloarchaeal RFO recovered from this search *(Trametes versicolor* EIW60452), a phylogeny was inferred for all 170 haloarchaeal opsins and the *T. versicolor* RFO. Sequences were aligned using Muscle 3.8.31 (Edgar, 2004b, 2004a) and alignment manually trimmed in Jalview (Clamp *et al*, 2004). After trimming, 220 positions were used in phylogenetic inference using FastTree (Price *et al*, 2010, 2009) and the resulting tree was visualized in FigTree (Rambaut, 2012). For tree see Fig. S4.

### Confirmation of native transcription of RFOs

Liquid cultures of *Hrr. litoreum* JCM 13561, *Hrr. distributum* JCM 9910, *Nab. magadii* DSM 3394 and *Nbt. gregoryi* SP2 were grown to mid-log phase in JCM 168 *(Hrr. litoreum* and *distributum)* or DSM 371 *(Nab. magadii* and *Nbt. gregoryi)* media. Cell pellets were collected from 2 mL of mid-log phase cultures and RNA harvested using 1.0 mL TRIzol^®^ Reagent with standard extraction protocol followed by a DNase digestion step and a second TRIzol^®^ extraction with 0.5 mL Trizol^®^ Reagent. DNase digestion was with NEB DNase I (M0303S) using standard protocol. Following second Trizol^®^ extraction, RNA was tested for gDNA contamination using PCR with haloarchaeal 16S primers (Table S3) prior to reverse-transcription. Reverse transcription was carried out with SuperScript^®^ III Reverse Transcriptase from Life Technologies™ using standard protocols, and random hexamers (Qiagen, 79236). Presence of transcript was confirmed via PCR with primers listed in Table S3. Cross-reactivity of primers for species with multiple RFO homologs was tested and primer sets found to be gene-specific. For *Hrr. distributum*, additional cultures were grown under the following conditions, and transcript presence verified using methods described above: a) 350 rpm, 3.42 M NaCl, b) 350 rpm, 5.0 M NaCl, c) 50 rpm, 3.42 M NaCl. Conditions (a) and (b) were incubated using a G-53 gyratory tier shaker (New Brunswick Scientific, M1074), condition (c) in an Innova 44R incubator (New Brunswick Scientific, M1282). For each condition, samples were collected at mid-log and stationary phase.

### Growth of *Halobacterium salinarum* clones

Unless otherwise specified, *Halobacterium salinarum* NRC-1 and derivatives were grown in CM medium (250 g/L NaCl, 20 g/L MgSO4 · 7H2O, 2 g/L KCl, 3 g/L Na3Citrate, 10 g/L Oxoid Neutralized Peptone (Oxoid, LP0037)) at 37°C in an Innova 44R incubator (New Brunswick Scientific, M1282) shaken at 175 rpm.

### Cloning

In order to heterologously express His-tagged RFO proteins, genes encoding *Nab. magadii* WP_004267173 and *Hrr. distributum* ELZ45759 were PCR amplified from genomic DNA using primers listed in Table S3 in 50 μL PCR reactions: 5 μL 10X buffer, 10 μL Q solution, 2.5 μL 10 μM forward primer, 2.5 μL 10 μM reverse primer, 1.25 μL 10 mM dNTPs, 0.3 μL TAQ polymerase (Qiagen, Q201203), 0.1 μL Pfu Polymerase (Stratagene, 600153), 28.35 μL MilliQ H_2_O using the following PCR cycle: 98°C 10 min, (98°C 30 sec, 55°C 1 min, 72°C 1 min 35 sec) x 25 cycles, 72°C 5 min in a Dyad Peltier Thermocycler (BioRad). Additionally, genes encoding a canonical BR (*Har. marismortui* YP_137573) and a colorless transmembrane protein *(Hbt. sp*. NRC-1 *pstC2* VNG0455G), were amplified as controls from genomic DNA as described above. Amplified genes were then ligated into two expression vectors, one under the control of the NRC-1 *bop* promoter (pDJLCHIS) and the other under the control of the NRC-1 ferredoxin promoter (pMTFCHIS2) [Data Dryad link] using the restriction enzymes NdeI (NEB, R0111S) and BamHI (NEB, R0136S for pMTFCHIS2) or HindIII (NEB, R0104S for pDJLCHIS). Clones were transformed into chemically competent DH5α cells and plated on LB agar plates containing 100 μg/mL carbenicillin (Fisher Scientific, BP26485). Success of cloning was verified through Sanger sequencing using plasmid specific sequencing primers (Table S3). Constructed pDJLCHIS and pMTFCHIS2 expression vectors (Fig. S6) were transformed into *Hbt. sp*. NRC-1 SD23 and *Hbt. sp*. NRC-1 SD20 strains, respectively.

Similarly, for heterologous expression in *Escherichia coli*, genes encoding *Nab. magadii* WP_004267173, *Hrr. distributum* ELZ45759, and *Har. marismortui* BR2 YP_137573 were PCR amplified from genomic DNA and cloned into pET29b+ (Novagen) using *NdeI* and *HindIII* restriction sites as described above. All cloned constructs, including an empty pET29b+ vector, were transformed into *E. coli* BLR(DE3) cells (Merck Millipore) for expression.

### Protein-level expression of RFOs in heterologous hosts

To express RFO proteins in *E. coli*, all clones were grown in Luria Broth (LB) under 50 μg/mL kanamycin resistance (kan) in an Innova 44R Shaking Incubator (New Brunswick) with shaking at 175 rpm. The pET29b+/RFO, pET29b+/BR2 and pET29b+/ empty BLR(DE3) clones were revived from freezer stock overnight in 4 mL of LB + kan at 37°C. Revived cells were re-cultured in 25 mL of LB + kan with a starting OD_600_ of 0.1 and grown at 37°C to mid-log (OD_600_ = 0.4). The mid-log subcultures were then used to inoculate 1 L LB + kan with a starting OD_600_ of 0.01 and placed in the 37°C shaking incubator. Once the cultures returned to mid-log (OD_600_ = 0.4), all cultures were induced with 0.5 mM IPTG (Fisher Scientific, BP1755), supplemented with 10 μM all-trans retinal (Sigma-Aldrich, R2500), and incubated at 18°C for 18 hours with shaking. Cells were harvested by centrifuging at 8000 rpm for 10 min and stored at -80°C until use.

To purify, cell pellets were thawed and re-suspended in 15 mL of solubilization buffer (100 mM Na/K phosphate pH 7.4, 2% Triton-X100, 10 μM all-trans retinal) with one cOmplete^®^ Mini EDTA free protease inhibitor cocktail tablet (Roche, 04693159001). Samples were sonicated three times with a Model 120 Sonic Dismembrator (Fisher Scientific, FB120) fitted with a Model CL-18 probe at 65% power alternatively for 2 seconds ON and 2 seconds OFF; for a total of 2 minutes ON. Sonicated samples were incubated on a rotisserie for 3 hours at 4°C. Cell debris was spun down by centrifugation at 8000 rpm for 10 minutes, and the supernatant was mixed with an equal volume of equilibration/wash buffer (50 mM Na+ phosphate, 300 mM NaCl, 10 mM imidazole; pH 7.4, 0.5% Triton-X100, 10 μM all-trans retinal). 250 μL of HisPur™ Cobalt Resin (Thermo Scientific, 89964) was washed with 500 μL of equilibration/wash buffer and mixed with equilibrated lysate for 45 minutes in a rotisserie at 4°C. Conjugated resin was washed three times with 500 μL equilibration/wash buffer and eluted twice with 250 μL of elution buffer (50 mM Na+ phosphate, 300 mM NaCl, 150 mM imidazole; pH 7.4, 0.5% Triton-X100, 10 μM all-trans retinal). 250 μL of the first eluent was concentrated to ~100 μL using a Microcon 30 kDa MWCO centrifugal filter column (Millipore, 42410) and aliquoted into a Greiner Half Area UV-Star^®^ microplate (Greiner Bio-One, 675801). Absorbance was measured from 250–800 nm in 2 nm intervals with an Infinite M200 plate reader (Tecan, 30016056). Background (elution buffer) was subtracted from all spectra and spectra were normalized to the 280 nm absorbance value of the *Har. marismortui* BR2 positive control. For spectra see Fig 4C.

To verify expression of RFO protein, we performed a Western blot on HisPur™ purified lysate from cloned RFOs in *E. coli* BLR(DE3) background using the following protocol: SDS-PAGE was run as described in SI Methods and the gel was soaked in 15 mL of transfer buffer (25 mM Tris base, 192 mM glycine, 10% methanol, pH 8.4) for 15 min. A 0.2 μM pore diameter nitrocellulose membrane (Biorad, 162–0112), foam pads, and 3 mm Whatman filters were soaked in transfer buffer before assembling the sandwich for transfer. Proteins were transferred from the PAGE gel to the nitrocellulose membrane at 30V for 1 hr on ice. His-tagged proteins were probed using the SuperSignal^®^ West HisProbe™ Kit (Thermo Scientific™, 15168) and signal was detected using Amersham ECL Prime Western Blotting Detection Reagent (GE Healthcare Life Sciences, RPN2232). For Western blot, see Fig. 4B.

### Protein-level expression of RFOs in native hosts

Several methods were used to investigate protein-level expression of RFO homologs in both native and heterologous hosts. First, we attempted identification of RFOs by LC-MS/MS in the native host *Nbt. gregoryi* SP2, which has four unique RFO genes. Cell pellets collected from mid-log phase cultures were lysed with 200 μL lysis buffer containing 100 mg SDS, 10 mL diH_2_O, 60 μL DNase I (GoldBio D-300–1), 0.25 mg RNase A (Roche, 10109169001), and 1 cOmplete^®^ Mini EDTA free protease inhibitor cocktail tablet (Roche, 04693159001) per 15 mL buffer. Lysate was sonicated for 20 minutes (30 s on/off) on a Biorupter^®^ UCD-200 (Diagenode). Cellular debris was removed by centrifugation, and supernatant denatured for 10 min at 100°C with 6x SDS buffer. Approximately 40 μg protein was loaded into pre-poured 4–20% SDS-PAGE gel (Bionexus Inc., 2BNPC420) and run approximately 1 cm into gel. Total protein band was excised and subjected to in-gel trypsin digest according to the following protocol: gel was cut into 1 mm^3^ pieces, washed with 50 mM Ammonium Bicarbonate (AmBic), shrunk with acetonitrile (ACN), reduced with 10 mM DTT/50 mM AmBic, shrunk again with ACN, incubated in 55 mM iodoacetamide/50mM AmBic 20 min in the dark, washed with 50 mM AmBic, shrunk with ACN and partially dried in a vacuum concentrator (Labconco). Overnight digestion was carried out at 37°C with 250 ng of trypsin (Promega, V5117) in 50 mM AmBic (pH 8). The supernatant was sonicated in 60% ACN and 0.1% trifluoroacetic acid for 10 min, then dried in the vacuum concentrator. Digested peptides were analyzed by LC-MS/MS on a Thermo Q-Exactive mass spectrometer with Michrom Paradigm LC and CTC Pal autosampler. Peptides were directly loaded onto an Agilent ZORBAX 300SB C18 reversed phase trap cartridge, which, after loading, was switched in-line with a Michrom Magic C18 AQ 200 um x 150 mm C18 column connected to a Thermo-Finnigan LTQ iontrap mass spectrometer through a Michrom Advance Plug and Play nano-spray source. The nano-LC column (Michrom 3μ 200Å MAGIC C18AQ 200Å x 150 mm) was used with a 90 min-long gradient (1–10% buffer B in 5 min, 10–35% buffer B in 65 min, 35–70% buffer B in 5 min, 70% buffer in 1 min, 1% buffer B in 14 min) at a flow rate of 2 uL min^-1^ for the maximum separation of tryptic peptides. A top 15 method was used with Xcalibur software to collect Q-Exactive data, with a scan range of 300–1600 m/z. Results were searched against a *Nbt. gregoryi* database with cRAP proteins and reversed sequences (5402 proteins total) in X!Tandem (The Global Proteome Machine Organization, 2013) with a fragment ion mass tolerance of 20 PPM and a parent ion tolerance of 20 PPM. Carbamidomethyl of cysteine was specified in X! Tandem as a fixed modification. Glu->pyro-Glu of the N-terminus, ammonia-loss of the N-terminus, gln->pyro-Glu of the N-terminus, deamidation of asparagine and glutamine, oxidation of methionine and tryptophan, dioxidation of methionine and tryptophan and acetylation of the N-terminus were specified in X!Tandem as variable modifications. Scaffold v.4.0.7 (Proteome Software Inc.) was used to validate peptide and protein identifications. Peptide identifications were accepted if they could be identified with confidence of greater than or equal to 95% and protein identifications were accepted if they could be identified with confidence of greater than or equal to 95% and contained at least two identified peptides. Protein probabilities were assigned by the Protein Proph*et al*gorithm (Nesvizhskii *et al*, 2003). Proteins that contained similar peptides and could not be differentiated were grouped. Proteins sharing significant peptide evidence were grouped into clusters. No RFO proteins were identified from *Nbt. gregoryi* SP2 lysate.

After failing to detect RFO protein in a native host, we purified His-tagged cloned proteins from a *Hbt. sp*. NRC-1 SD23 *bop*(-) background using HisPur^®^ Cobalt resin (Thermo Scientific™, 89965). Exudate was loaded onto a 4–20% SDS-PAGE gel as described above, and size separated. Based upon presence of band in the three experimental samples and absence in *Hbt. sp*. NRC-1 SD23 control lacking expression vector, the region of the gel corresponding to 50 kDa MW proteins was excised and prepared for proteomics as described above. No RFO proteins were detected.

We next performed Western blot on lysate from cloned RFOs in *Hbt. sp*. NRC-1 SD23 bop(-) background using the following protocol. SDS-PAGE was run as described above and gel soaked in 15 mL of transfer buffer (25 mM Tris base, 192 mM glycine, 10% methanol, pH 8.4) for 15 min. PVDF membrane (Novex, LC2002) was successively soaked in 100% methanol, MilliQ™ H2O, and transfer buffer. Foam pads and 3mm

Whatman filters were also soaked in transfer buffer. Western gel was run at 30V for 1 hr. His-tagged proteins were probed using the SuperSignal^®^ West HisProbe™ Kit (Thermo Scientific™, 15168). Based upon presence of two bands at ~15 and ~20 kDa in the three experimental samples, the region of the gel corresponding to 13–25 kDa was extracted and prepared for proteomics as described above. No RFOs were detected, however, the three opsin proteins present in the host (HR, SR1, and SR2) were observed, indicating the region of the gel used was the correct MW range for opsins. Over 900 proteins were identified with a protein false discovery rate of 1.7 *%* and peptide false discovery rate of 0.36%, using a *Hbt. sp*. NRC-1 database with RFOs added.

## Data Access

Haloarchaeal genomes can be accessed through the NCBI Genomes database under the following accession numbers: AOHS-AOIE, AOII-AOIW, AOJD-AOJO, AOLD-AOLS, AOLW-AOMF.

## Acknowledgments

The authors would like to thank past and current members of the Facciotti and Eisen labs for many productive discussions about our “odd opsins”.

## Author Contributions

Designed and planned the study: EAB, MTF. Performed the experiments: EAB, AIY, PMS, TK, TW, RE, VY. Analyzed the data: EAB, AIY, MTF, PMS, TK, KSYS, VY. Wrote the manuscript: EAB, MTF, VY, TK, AY.

## Disclosure Declaration

PMS is currently an employee of Proteome Software, Portland, Oregon, USA. However, this employment began after completion of work on this manuscript. The authors declare that no competing interests exist.

